# No evidence of musical training influencing the cortical contribution to the speech-FFR and its modulation through selective attention

**DOI:** 10.1101/2024.07.25.605057

**Authors:** Jasmin Riegel, Alina Schüller, Tobias Reichenbach

## Abstract

Musicians can have better abilities to understand speech in adverse conditions such as background noise than non-musicians. However, the neural mechanisms behind such enhanced behavioral performances remain largely unclear. Studies have found that the subcortical frequency-following response to the fundamental frequency of speech and its higher harmonics (speech-FFR) may be involved since it is larger in people with musical training than in those without. Recent research has shown that the speech-FFR consists of a cortical contribution in addition to the subcortical sources. Both the subcortical and the cortical contribution are modulated by selective attention to one of two competing speakers. However, it is unknown whether the strength of the cortical contribution to the speech-FFR, or its attention modulation, is influenced by musical training. Here we investigate these issues through magnetoencephalographic (MEG) recordings of 52 subjects (18 musicians, 25 non-musicians, and 9 neutral participants) listening to two competing male speakers while selectively attending one of them. The speech-in-noise comprehension abilities of the participants were not assessed. We find that musicians and non-musicians display comparable cortical speech-FFRs and additionally exhibit similar subject-to-subject variability in the response. Furthermore, we also do not observe a difference in the modulation of the neural response through selective attention between musicians and non-musicians. Moreover, when assessing whether the cortical speech-FFRs are influenced by particular aspects of musical training, no significant effects emerged. Taken together, we did not find any effect of musical training on the cortical speech-FFR.

**Significance statement:** In previous research musicians have been found to exhibit larger subcortical responses to the pitch of a speaker than non-musicians. These larger responses may reflect enhanced pitch processing due to musical training and may explain why musicians tend to understand speech better in noisy environments than people without musical training. However, higher-level cortical responses to the pitch of a voice exist as well and are influenced by attention. We show here that, unlike the subcortical responses, the cortical activities do not differ between musicians and non-musicians. The attentional effects are not influenced by musical training. Our results suggest that, unlike the subcortical response, the cortical response to pitch is not shaped by musical training.

## Introduction

Speech comprehension is essential for human interaction and communication, yet the underlying neural mechanisms remain incompletely understood. Hearing-impaired people often face major difficulty with understanding speech in noisy environments, including many social settings such as pubs or restaurants (McDermott, 2009; Bronkhorst, 2000). Some studies showed that people with extensive musical training can be particularly apt at understanding speech in background noise (Parbery-Clark et al., 2009, 2012a). This is especially the case in difficult listening conditions. A review and meta-analysis found that especially for background noise with more than one speaker musicians tend to perform better than non-musicians (Maillard et al., 2023). It is important to mention, though, that this has not been found in every study. In fact, out of seven investigations which all had two speakers as disturbing background noise, four studies found enhanced performance by musicians (Deroche et al., 2017; Kaplan et al., 2021; Morse-Fortier et al., 2017; Zhang et al., 2020), whereas three did not find differences (Deroche et al., 2017; Madsen et al., 2019; Zhang et al., 2020).

Despite this variety in the obtained results, a survey of many such studies described a positive impact of musical training on the understanding of speech in noise (Maillard et al., 2023). Investigating the neural mechanisms that lead to this improved behavioral performance may shed light both on the effects of musical training as well as on mechanisms for speech-in-noise comprehension.

Several characteristics of speech may aid musicians in their enhanced speech-in-noise comprehension. Pitch, timbre, and timing are, for instance, important attributes of speech and also matter for the processing of musical signals (Magne et al., 2006; Besson et al., 2007; Musacchia et al., 2007). An investigation from Pantev et al. (2001) showed indeed that musicians had stronger timbre-specific responses in the auditory cortex when presented with tones of their trained instruments than non-musicians. Furthermore, musically trained subjects were also observed to exhibit stronger auditory-evoked cortical responses to signals presented in high levels of noise (Meha-Bettison et al., 2017).

Pitch is an integral part of most music, and some studies accordingly found musicians to be better in pitch perception and discrimination than non-musicians (Magne et al., 2006; Bianchi et al., 2016; Toh et al., 2023). The enhanced ability of musicians to discriminate speech in noise may therefore also involve a better neural representation of the pitch of a voice.

The sensation of a voice’s pitch results from its temporal fine structure. The latter consists of a fundamental frequency, arising from the rhythmic opening and closing of the glottis, and its higher harmonics. The temporal fine structure of a voice can provide a cue that can aid a listener to focus on a target voice amidst background noise such as other competing speakers (Hopkins and Moore, 2009; Eaves et al., 2011).

The temporal fine structure of speech elicits a neural response at the fundamental frequency as well as, to a lesser degree, at the higher harmonics (Russo et al., 2004; Skoe and Kraus, 2010). Because the response at the fundamental frequency is reminiscent of the frequency-following response (FFR) to a pure tone, we will in the following refer to it as speech-FFR. The speech-FFR can be measured non-invasively through electroencephalography (EEG) or magnetoencephalography (MEG). While subcortical sources of the speech-FFR in different parts of the brainstem and midbrain are well established, recent MEG measurements have also revealed contributions from the auditory cortex (Coffey et al., 2016, 2019; Kulasingham et al., 2020).

Following the aforementioned hypothesis that musicians may have an enhanced representation of pitch, the subcortical contributions to the speech-FFR were found to be higher in people with musical training as compared to those without (Musacchia et al., 2007; Parbery-Clark et al., 2012a). Musicians also exhibited more robust subcortical representations of acoustic stimuli in extremely noisy environments (Parbery-Clark et al., 2012b). In line with these findings, native speakers of Mandarin, a tonal language in which pitch can differentiate between different words, were found to have enhanced brainstem contributions to FFRs as compared to native speakers of English, which is not tonal (Bidelman et al., 2011).

Both the subcortical and the cortical contribution to the speech-FFR are modulated by selective attention to one of two competing speakers. The attentional modulation has first been shown for the subcortical signals using EEG (Forte et al., 2017; Etard et al., 2019; Saiz-Aĺıa et al., 2019), and more recently for cortical responses through MEG (Schüller et al., 2023a; Commuri et al., 2023).

However, it has not yet been investigated whether the cortical contribution to the speech-FFR is influenced by musical training, nor whether musical training affects attentional modulation. This study aims to address this gap in knowledge by recording MEG responses to two competing voices from participants with different amounts of musical training.

## Materials and methods

### Experimental design and data analysis

The participants were presented with audio signals from two competing speakers and asked to attend one at a time while their MEG was recorded (Figure 1). The resulting signal was source reconstructed. The signals at the sources in the auditory cortex were related to two features of one of the voices through ridge regression, to capture different aspects of the speech-FFR. We thus obtained temporal response functions (TRFs) that described the speech-FFRs. The TRFs were subsequently compared between musicians and non-musicians for both attending and ignoring the speaker.

### Participants

The study included 52 healthy participants aged between 18 and 30 years (musicians: male: 8, female: 10; non-musicians: male: 13, female: 12; neutral: male: 5, female: 4). All participants were right-handed, had no history of hearing impairments or neurological diseases, had no metal objects inside their bodies, and were native German speakers. They were recruited in Erlangen, Germany.

Out of the 52 subjects, 16 had participated in a previous study with the same setup conducted in our group (Schüller et al., 2023a). To be assigned to the two groups, “musicians” and “non-musicians”, the subjects had to meet specific criteria, based on their responses to a questionnaire on musical training, which is presented in detail later.

Participants were categorized as musicians if they had started playing an instrument at the age of seven or earlier, had played instruments for a total of ten or more years, and were currently playing an instrument. The choice of instruments was flexible, but drums were not considered.

Participants were categorized as non-musicians if they had never played an instrument or had played an instrument for a maximum of three years, and if so, they had started playing the instrument at the age of seven or older.

All subjects of the data set of the previous study were retrospectively asked to fill out the questionnaire on musical training. Among them were five musicians, two non-musicians, and nine subjects who did not belong to either group. With the addition of the newly recruited subjects, our study consisted of 18 musicians, 25 non-musicians, and nine individuals who were not categorized into either group and were considered neutral. Although the study was unbalanced with respect to the two groups of musicians and non-musicians, each group was balanced with respect to gender.

**Figure 1:**
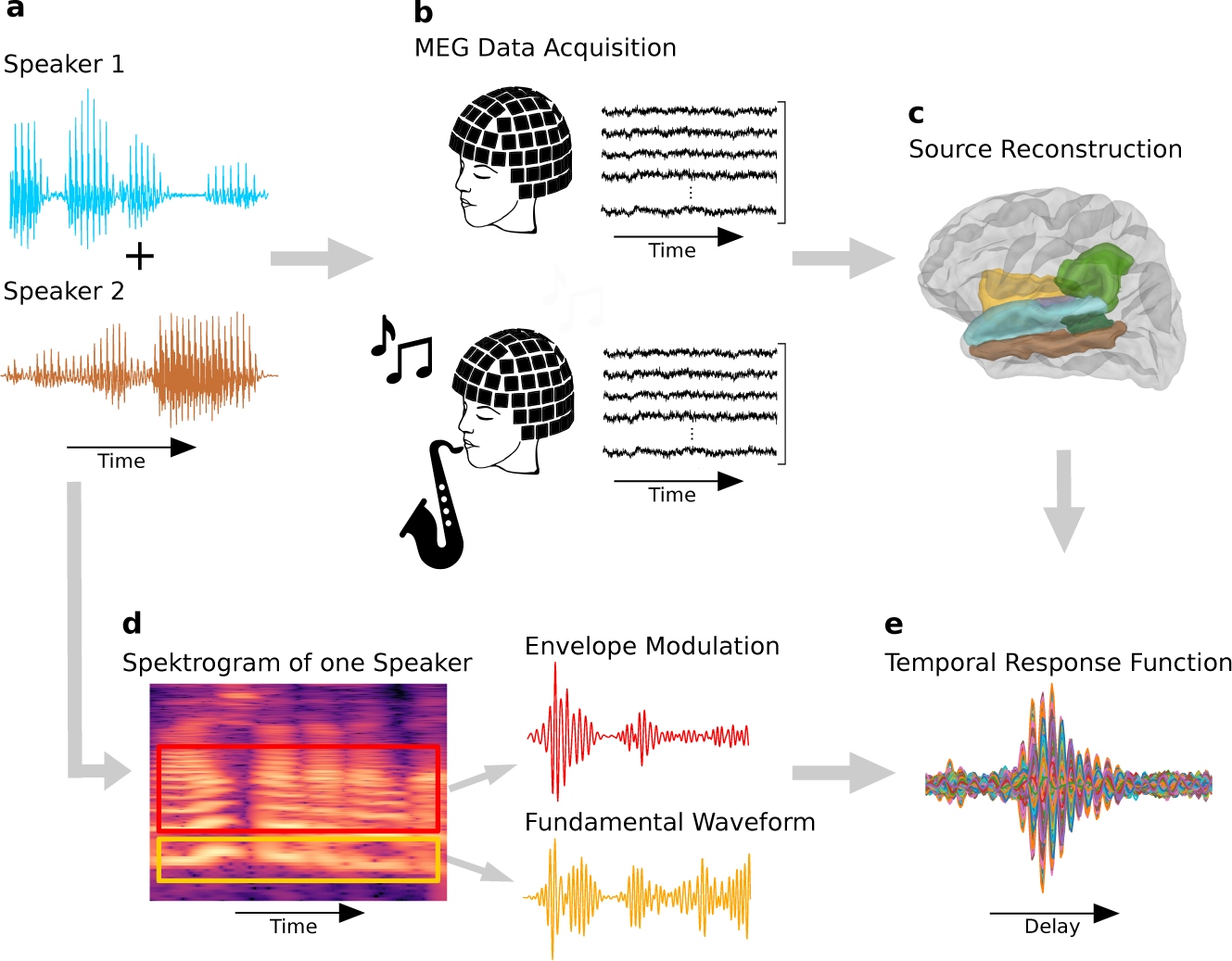
Experimental setup and acoustic stimuli. (a) Two audiobooks (one attended, and one ignored) were presented simultaneously while MEG was recorded. (b) MEG recordings of participants with different amounts of musical training were conducted. (c) The MEG recordings were source reconstructed to obtain their origin in a cortical region of interest (purple: transverse temporal gyri, blue: superior temporal gyri, brown: middle temporal gyri, dark green: banks of the superior temporal sulci, yellow: supramarginal gyri and light green: insular cortex). (d) The spectrogram of the acoustic input signal was analyzed and processed into two features. The fundamental waveform was obtained by band-pass filtering the signal around the fundamental frequency (yellow). The higher-mode envelope modulation reflected the amplitude modulations of the higher harmonics (red). (e) Temporal Response Functions (TRF) depicting the speech-FFRs are generated for each participant by processing the two speech features and the acquired MEG recordings in a linear forward model.

### Measuring procedure and acoustic stimuli

During the measurement, the participants listened to acoustic stimuli consisting of two competing audio files diotically, both narrated by male speakers.

A total of four audiobooks were used in the study. Two of these audiobooks (”story audiobooks”) were those that the participants were asked to attend, while the other two were used as distractor signals (”noise audiobooks”). The first story audiobook was “Frau Ella” by Florian Beckerhoff, and the first noise audio-book was “Darum” by Daniel Glattauer. Both of these books were narrated by Peter Jordan. The second story audiobook was “Den Hund überleben” by Stefan Hornbach, and as the second noise audiobook we employed “Looking for hope” by Colleen Hoover (translated to German by Katarina Ganslandt). Both of these books were narrated by Pascal Houdus. All four audiobooks were published by Hörbuch Hamburg.

The first speaker, Peter Jordan, is referred to as the LP speaker due to his voice having, on average, a lower pitch compared to Pascal Houdus, who is referred to as the HP speaker. Peter Jordan’s fundamental frequency had a frequency range of approximately 70 - 120 Hz, while Pascal Houdus’s fundamental frequency varied between approximately 100 - 150 Hz.

The first and second audiobook were presented in an alternating manner. The first story audiobook was accompanied by the second noise audiobook. Conversely, the second story audiobook was presented in combination with the first noise audiobook as the distractor signal. The participants were informed on which book to attend by letting the target voice begin speaking five seconds before the distracting speaker. Each participant listened to ten chapters, resulting in approximately 40 minutes of audio sequences in total.

The sound-pressure level of the stimuli was *∼*67 dB(A) during the experiment for both the attended and ignored voices. To make the listening process easier for the participants, the original chapter lengths of the attended audiobooks were used. As a result, the chapters had varying lengths between three and five minutes. For the ignored audiobooks, random parts of the audio files were selected to match the expected lengths.

To avoid eye movement artifacts during the measurement, participants were asked to focus on a fixation cross shown on a display above their heads. After each chapter, three single-choice questions with four possible answers appeared on the screen consecutively, and the participants were asked to provide the correct answers. This allowed to verify whether they were listening to the correct speaker.

Although in this work only MEG data was analyzed, we also measured the electroencephalogram (EEG), the electrocardiogram, and the electrooculogram simultaneously.

### Experimental setup

The setup used for the presentation of the speech stimuli was already prepared through previous studies. Initially, it was built for a study by Schilling et al. (2021). It consisted of two computers, one for stimulation and one for recording. The stimulation computer was connected to a USB sound card which provided five analog outputs.

The first two outputs were connected to an audio amplifier (AIWA, XA-003) which was connected via audio cables to two speakers (Pioneer, TS-G1020F), each creating the signal presented to one of the subject’s ears. From each speaker, a silicone funnel transferred the sound wave into flexible tubes of approximately 2 m length and 2 cm inner diameter. These tubes were the only part of the auditory setup entering the magnetically shielded chamber in which the MEG was situated to protect it from environmental noise. The tubes entered the chamber via a small hole. Because of the length of the tubes a constant time delay was created between the arrival of the sound in the subject’s ear and the earlier generation. This delay was 6 ms and was taken into account for the alignment between the MEG data and the sound stimuli.

The first output of the audio amplifier was connected in parallel to an analog input channel of the MEG data logger. This allowed for the recorded MEG signals and the stimuli to be aligned with a precision of 1 ms.

To prevent temporal jittering of the signal due to, for instance, multi-threading of the stimulation PC’s operating system, a forced alignment of the presented signal and the MEG recording was created. This was facilitated via the third analog output of the sound device which was utilized to transmit trigger pulses derived from the forced alignment process to the MEG recording system through an additional analog input channel.

### Data acquisition and preprocessing of MEG data

The measurement took place at the University Hospital in Erlangen, Germany. A 248 magnetometer system (4D Neuroimaging, San Diego, CA, USA) was utilized. The subjects were instructed to lie down with their head positioned in the MEG system, which sampled at a frequency of 1,017.25 Hz. Before commencing the measurement, the head shape of each subject was digitized, and five landmark positions were recorded.

Throughout the measurement, an analog band-pass filter with a range of 1.0 *−* 200 Hz was employed to process the signals. Additionally, the system featured a calibrated linear weighting applied to 23 reference sensors, developed by 4D Neuroimaging in San Diego, CA, USA. These reference sensors allowed to correct for environmental noise during the measurement process, enhancing the overall accuracy and reliability of the collected data.

After the measurement, several further filters were applied to the acquired data. First, a 50 Hz notch filter (firwin, transition bandwidth 0.5 Hz) was applied, a standard procedure for removing power line interference. Second, the data was down-sampled to 1,000 Hz. Third, another band-pass filter was applied to select the range of the fundamental frequency of the LP speaker. A previous study with the same setup indeed found that the cortical speech-FFRs evoked by the LP speaker were much stronger than those by the HP speaker (Schüller et al., 2023a). This is in line with previous findings that speech-FFRs are stronger at lower fundamental frequencies (Saiz-Alía et al., 2019; Saiz-Alía and Reichenbach, 2020; Van Canneyt et al., 2021). Here we accordingly only analyzed the neural responses to the LP speaker, using band-pass filter borders between 70 - 120 Hz. The band-pass filter was a linear digital Butterworth filter of second order with a critical frequency obtained by dividing the lower and upper cut-off frequency by the Nyquist frequency, applied forward and backward to prevent phase delays.

### Processing of the acoustic stimuli

In line with previous publications, we employed two speech features to obtain the speech-FFRs (Kulasingham et al., 2020; Kegler et al., 2022a; Schüller et al., 2023a). The first one was the fundamental waveform *w_t_* and the second one the envelope modulation *e_t_* of the higher modes of the fundamental frequency of the speech signals.

The Probabilistic YIN (pYIN) algorithm (Mauch and Dixon, 2014) was employed to extract the fundamental frequency of a speech signal. We then bandpass-filtered the speech signal of the LP speaker in the range of the corresponding fundamental frequency, between 65 Hz and 120 Hz (IIR-filter applied forward-backward, 4th order), yielding the fundamental waveform.

To obtain the second speech feature, the envelope modulation, we first processed the speech signal of the LP speaker by a model of the auditory periphery, encompassing the tonotopic properties of the cochlea, the frequency tuning of auditory-nerve fibers, and the neural responses observed in various brainstem nuclei. We thus used a translation of the original Matlab code (Chi et al., 2005) into Python.

The model utilizes a series of constant-*Q* bandpass filters, which are specialized filters designed to have varying frequency resolution throughout the auditory spectrum. These filters are subsequently followed by nonlinear compression and derivative mechanisms operating across different scales, resulting in enhanced frequency resolution. Each frequency band undergoes an envelope detection process that extracts the signal’s amplitude envelope at that specific frequency. The resulting envelope modulation is then band-pass filtered in the range of the fundamental frequency of the LP speaker (70 - 120 Hz, IIR-filter applied forward-backward, 4th order).

To assess to which degree the two signals captured different aspects of the speech stimulus, we computed Pearson’s correlation coefficient. We found that the correlation was very low but highly statistically significant (*r* = 0.009 and *p* = 8*e −* 26). The low value of the correlation coefficient near zero evidenced that the two speech features were essentially unrelated to each other and thus represented different aspects of the signal. The high statistical significance of the very small correlation value reflects the lack of noise in the speech signal from which the two features were obtained.

### Source reconstruction

Source reconstruction aims to identify the locations that generated the measured signal. By applying a forward model to a predefined source space, the relationship between the possible source points and the resulting measured signal was determined. This resulted in an individual leadfield matrix for every subject. Subsequently, through the application of a beamformer, the source reconstructed MEG data was generated.

For an individual calculation of the source reconstruction for each participant, further information was required which was provided by the digitized head shape that was measured for every subject. Firstly, coregistration between the position of the sensors on the scalp and the MEG sensors was carried out. The spatial relationship of each subject’s head with the MEG scanner was thus determined using five marker coils. Their positions were recorded at the beginning and end of the measurement.

Secondly, the anatomy of the brain is required as it defines the possible source points that generate the signal. Ideally, a magnetic resonance imaging (MRI) scan for each participant would have been available for individual source reconstruction. However, previous studies have explored the substitution of MRI scans with an average brain template, yielding comparable results (Douw et al., 2018; Holliday et al., 2003; Schüller et al., 2023b; Kulasingham et al., 2020). Therefore, a template provided by Freesurfer (Fischl, 2012) was used to create a target volume through rotation, translation, and uniform scaling of an average brain volume named ‘fsaverage’ to match the digitized head shape as well as possible. This entire analysis was carried out using the MNE-Python software package (Gramfort et al., 2014).

Previous research showed that the cortical contributions originate in the left and right auditory cortex (Schüller et al., 2023b). We, therefore, considered a region of interest (ROI) that contained the transverse temporal, middle temporal, superior temporal, and supramarginal gyri, as well as the insular cortex and the banks of the superior temporal sulci (Figure 1 (c)). The so-defined ROI contained 525 source points. They were utilized to create a regular grid with a spacing of 5 mm, thereby producing a volumetric source space. The chosen volume is presented in Figure 1 (c). For each of the source points, we computed a TRF curve. The magnitudes of the resulting 525 TRF curves were then averaged for each participant. The resulting value at the peak time lag was then used for further analysis in Figure 2 - 4.

With the possible source points set up, the next step was to run the forward model generating the leadfield matrix. The forward model individually calculates the influence of a signal generated at every single source point in the source space onto all possible measurement points. It thereby utilizes the individual sensor locations, regions of interest in the generated brain volume, and the data covariance and noise covariance matrix. The latter were generated by using MEG data from a one-minute interval during which the acoustic stimulus was presented, and by recording MEG data from a three-minute pre-stimulus empty room phase, respectively. The resulting leadfield matrix contained a row for each MEG sensor and a column for each source point, describing their relationship in terms of dipole orientation and signal strength.

For the final step of source reconstruction, the linearly constrained minimum variance (LCMV) beam-former (Bourgeois and Minker, 2009) was computed on the leadfield matrix and the individual volume source space. The beamformer was constructed in a way that enhanced the activity of source points and reduced the signal from interfering sources. At every source point, we thus obtained an estimation of the current dipole in the form of a three-dimensional vector.

### Derivation of temporal response functions (TRFs)

We determined the speech-FFRs by computing Temporal Response Functions (TRFs). The TRFs are the coefficients of a linear forward model that reconstructed the MEG data from the two speech features, the fundamental waveform *f_t_* and the envelope modulation *e_t_*. The linear forward model thereby employed different latency shifts *τ* of the acoustic features with respect to the MEG data, so that neural responses at different delays could be assessed. The neural response 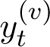 at time *t* in source voxel *v* was thus estimated as

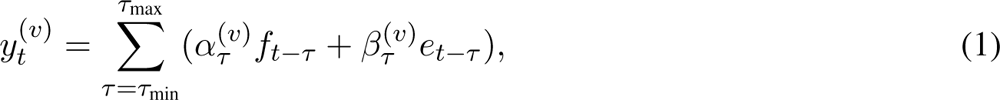

in which the coefficients 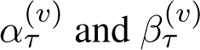 represent the TRFs.

To prevent overfitting, and in line with previous studies from the same research group as well as others, we used regularized ridge regression to compute the forward model (Wong et al., 2018). We thereby observed that the regularization parameter *λ* = 1 yielded optimal or near-optimal results for all subjects, and thus employed this value.

We considered latencies between a minimal delay of *τ*_min_ = *−*20 ms and a maximal delay of *τ*_max_ = 120 ms, with an increment between neighboring latencies of 1 ms, since the sampling rate was 1, 000 Hz. The TRFs were computed for each subject for all source points in the region of interest. The subject-specific TRFs were then averaged across subjects to obtain results on the population level.

Because the TRFs describe the relation of the fundamental waveform respectively the envelope modulation at the fundamental frequency to the MEG data, they display oscillations around the fundamental frequency. This oscillatory behavior was still visible in the amplitudes of the TRFs. To obtain a smoother shape of the magnitudes that is more amenable to subsequent analysis, we also computed the envelope of the magnitude of the TRFs.

This envelope was determined by first computing the Hilbert transform of the amplitude of a TRF. We then determined the absolute values and applied a Butterworth zero-delay low-pass filter (cut-off frequency = 70 Hz, order = 5), resulting in an envelope representation of the TRFs. The calculation of the envelope was done for each subject, and from there the population average was computed.

### Categorization of musicians and non-musicians

We employed three different criteria to categorize a subject as a musician. The first factor was the subject’s starting age of musical training, as has been established in various studies. For instance, research on neural activity while perceiving piano notes indicated a correlation between activation and the age at which piano training commenced (Pantev et al., 1998). Additionally, studies exploring speech comprehension in noisy environments revealed that musicians exhibit enhanced speech processing skills, which are further enhanced by early-life music training (Parbery-Clark et al., 2009). Similarly, in a study by Kraus and Chandrasekaran (2010), the age of training onset was identified as a determinant of neural plasticity associated with musical training. A study by Watanabe et al. (2006) examining the impact of early musical training on adult motor performance found that musicians commencing training before the age of seven demonstrated enhanced sensory-motor integration. Following these findings and other related ones, we employed a threshold of seven years for the starting age.

As the second criterion, we employed the total number of years the subject had trained in his or her life so far, at a certain minimal amount per week. Indeed, Kraus and Chandrasekaran (2010) showed in their study on neural plasticity that the number of years of continuous training had a relevant influence. Parbery-Clark et al. (2009) measured an advantage in processing speech in challenging listening environments due to sensory abilities that develop over the lifetime through consistent practice routines. Also, a study from Strait et al. (2009) stated that the magnitude of their measured neural responses to a complex auditory stimulus correlated with the number of years of consistent musical practice. The duration of musical training was thereby set to ten years, with at least three hourly training sessions per week and additional music lessons (Strait et al., 2009). Here, we similarly set the threshold for the total amount of musical training to ten years but only specified a training amount of at least an hour per week.

The third criterion was the number of years since their last training period, for which we set the threshold to zero. This criterion ensured that participants classified as musicians were currently playing an instrument. The criterion was adopted since it appeared as a criterion for non-musicians in several previous studies (Wong et al., 2007; Musacchia et al., 2007; Strait et al., 2009; Parbery-Clark et al., 2009,?; van Zuijen et al., 2005; Perrot et al., 1999).

In most studies examining musicians, any participant who is not categorized as a musician is denoted a non-musician. We employ a stricter approach here. We only categorize a subject as a non-musician if they haven’t had musical training before the age of seven, are presently not training, and if they had previously it lasted at most three years.

The criteria employed for categorizing subjects as musicians or non-musicians are summarized in Table 1.

**Table 1:** Criteria of musicians and non-musicians.

|  | Starting age | Total years of training | Presently training |
| --- | --- | --- | --- |
| Musicians | $\leq 7$ | $\geq 10$ | yes |
| Non-musicians | $\geq 7$ | $\leq 3$ | no |

### Scores of musical training

In addition to categorizing participants as musicians, non-musicians, or as neither of these two groups we also considered how the speech-FFRs depended on four different scores of musical training. The first two scores were the first two criteria for musicians: the starting age of training and the total duration of training. These two properties have also been taken as scores in the previously mentioned study about speech processing in challenging listening environments (Strait et al., 2009).

As a third score, we employed the momentary amount of training, since this measure reflects the current commitment of the participant to his or her musical training. The fourth score was the number of instruments the subject had been playing over their lifetime. We then also computed an average score of musical training. To this end, the four scores were normalized with respect to their maximal value and then added up. The four scores as well as the average score were calculated for all subjects, i.e. musicians, non-musicians, and participants who fell between these two categories.

### Statistical significance of individual cortical speech-FFRs

Before further analyzing the neural responses, their statistical significance was tested. Therefore, as conducted in earlier studies, noise models were generated by reversing the audio features in time and then computing noise TRFs (Schüller et al., 2023b; Kegler et al., 2022b).

The TRFs of each subject and the noise TRFs were tested for a statistically significant difference by applying a bootstrapping approach with 10, 000 permutations over the subject’s noise models. This generated a distribution of noise model magnitudes over time lags. Then, for each time delay, an empirical *p*-value was estimated by computing the proportion of noise values exceeding the actual TRF magnitude. Finally, the Bonferroni method was applied to the *p*-values to account for multiple comparisons across the different time lags.

### Statistical significance of the attentional modulation

To assess the significance of the difference in the speech-FFRs when attending and when ignoring a speaker, we compared the envelopes of the TRFs. To this end, we determined for each participant and for each of the four conditions (fundamental waveform attended and ignored, envelope modulation attended and ignored) if there was a significant peak in the envelope of the corresponding TRF. We then compared the latencies of these peaks between the attended and the ignored condition across the subjects through Mann-Whitney-U tests. We used the Mann-Whitney-U test as the distributions were not normally distributed (Shapiro-Wilk test).

The magnitude of each subject’s envelope TRF was then calculated at the obtained time delay. This was done in the attended and ignored conditions and for each of the acoustic features. To select the appropriate statistical test to assess differences between the amplitudes, we first assessed that the distributions were not normally distributed (Shapiro-Wilk test). We then compared the amplitudes between the attended and the ignored condition through a Mann-Whitney-U test, one for each of the two speech features.

### Statistical analysis of the influence of musical training

The influence of musical training was assessed by considering the envelopes of the TRFs at the latency at which the envelope peaked on the population level. This latency was computed for all participants excluding outliers and the participants with insignificant peaks. Outliers were identified on the basis of the interquartile range and excluded to make sure the selected time lag was not distorted by extreme values. Specifically, if a participant’s peak latency was below *Q*1 *−* 1.5 *· IQR* or above *Q*3 + 1.5 *· IQR*, it was considered an ourlier.

*Q* hereby stands for quartile, and *IQR* denotes the interquartile range. Participants whose neural response did not exhibit a significant peak were also excluded since the absence of a peak meant that no peak latency could be determined. The participants with insignificant peaks were only excluded for the computation of the time lag itself and then rejoined for the analysis of the TRF values at the obtained time lag.

In the following analysis, we considered the magnitudes of the neural responses at the peak of the responses on the population level, that is, at the same latency for each participant. We included the participants that did not show significant peaks in their neural responses and only excluded data points that were identified as outliers based on the interquartile range.

We first compared the envelopes of the TRFs between the musicians and the non-musicians. The normality of the distribution was assessed through the Shapiro-Wilk test. If the two distributions that we aimed to compare were both normally distributed, we utilized a Student’s t-test, and otherwise a Mann-Whitney-U test. We then assessed whether the attentional modulation differed between musicians and non-musicians. Therefore, for each subject, we defined an attentional modulation score *Q* as the difference between the envelope TRF in the attended condition (*a*) and in the ignored condition (*i*), divided by the sum of the two values:

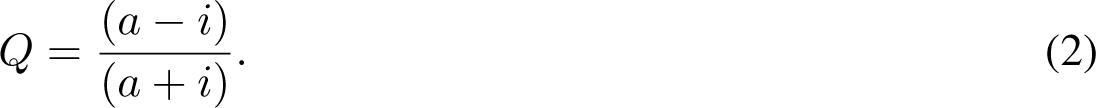

We then compared the scores *Q* between musicians and non-musicians through both an unpaired t-test and a Mann-Whitney-U test, as the Shapiro-Wilk test showed that the scores regarding the fundamental waveform followed a normal distribution, but not those for the envelope modulation.

The variances in the different measures (envelope of the TRFs at the latency of the maximum in the population average and the attentional modulation score) were assessed as well. In particular, we determined if the variances differed between musicians and non-musicians. This investigation was performed as, by visual inspection, the variances of the musicians and non-musicians seemed to differ. Furthermore, a larger variance for non-musicians might arise since they might differ in certain aspects of musical exposure, such as singing privately or extensive listening to music, that we did not assess. We, therefore, performed the Brown-Forsythe test for all distributions, as it is robust to departures from normality (Santos et al., 2018).

Next, we investigated whether the speech-FFRs depended on the scores of musical training. We, therefore, computed the Spearman correlation coefficient between the envelope of the TRFs and each of the different scores that described musical training (Kaptein and van den Heuvel, 2022). We then tested the obtained correlation coefficients for statistical significance. As this resulted in five different tests on the same data, we corrected for multiple comparisons through the false discovery rate from Benjamini Hochberg (Benjamini and Hochberg, 1995).

As a final investigation, we examined a possible correlation between the number of correct answers and the neural signal strength. Again the groups were not normally distributed and the correlation was therefore quantified through the Spearman correlation coefficient.

### Data and Code Accessibility

The MEG data is available at zenodo.org (https://zenodo.org/records/12793944). Python code for TRF analysis as described in this paper can be found on github.com (https://github.com/Al2606/MEG-Analysis-Pipeline).

## Results

### Attentional modulation of the cortical contribution to the speech-FFR

We first ascertained that the previously observed attention modulation of the cortical speech-FFR could be seen in our data as well. We, therefore, compared the TRFs for the fundamental waveform and for the envelope modulation between the attended and the ignored condition on the population level (Figure 2). Significant neural responses emerged at delays between about 20 ms and 50 ms (Figure 2 a - d). The envelopes of the TRFs amplitudes peaked at delays between 30 ms and 36 ms.

We wondered if there was a significant difference in the latencies at which the envelopes of the TRF amplitudes for individual subjects peaked, between the attended and the ignored condition. We thus compared the latencies of the peak responses of all subjects excluding outliers for each mode separately. In addition to the outliers also all participants with insignificant peaks were excluded. This resulted in excluding seven subjects for the attended and 18 for the ignored mode evaluated for the fundamental waveform. Furthermore for the envelope modulation when attending the speaker four subjects were excluded and when ignoring 22. The data from the included subject is shown as gray dots in Figure 2 (e),(f), together with the resulting box plots.

We used the Shapiro-Wilk test to define the type of distribution of our groups. As we had not normally distributed data, we applied the Mann-Withney-U test and observed no significant differences in the peak latencies between the attended and the ignored condition of the fundamental waveform and the envelope modulation (Figure 2 (e, f); Mann-Withney-U test; *p* = 0.06, fundamental waveform; *p* = 0.34, envelope modulation). As the mean latencies nonetheless differed in their absolute values, we used separate latencies for attended and ignored cases and for both fundamental frequency and envelope modulation. These latencies are those obtained in Figure 2 (e, f) including only subjects showing significant peaks. These latencies were 36 ms (fundamental waveform, attended), 39 ms (fundamental waveform, ignored), 32 ms (envelope modulation, attended), 36 ms (envelope modulation ignored).

Importantly, we found that the magnitude of the envelope at the peak latency was significantly higher when the speaker was attended than when he was ignored (Figure 2 (g,h); Mann-Withney-u test; *p* = 9.4*e −* 6, fundamental waveform; *p* = 7.4*e −* 5, envelope modulation). The Mann-Withney-U test was applied as the data were not normally distributed (Shapiro-Wilk test). The significant difference confirms the previously observed attentional modulation of the cortical speech-FFR (Schüller et al., 2023a; Commuri et al., 2023).

### No differences in the cortical speech-FFR between musicians and non-musicians

As a next step, we investigated whether the cortical speech-FFRs differed between the group of musicians and the group of non-musicians. We therefore computed the envelopes of the TRF magnitudes separately for the two groups, both in the attended and the ignored condition (Figure 3 (a)-(b)). As shown for the whole cohort in Figure 2, an influence of attention can also be detected for each of the two individual groups.

To assess potential differences between the two groups, we then focused again on the envelope at the peak latency (Figure 3 (c,d) and (e,f)). For each plot, the outliers were excluded separately. For the group of the non-musicians, this resulted in excluding one subject when attending and five when ignoring the speaker, regarding the response to the fundamental waveform. Furthermore, no subject was excluded when attending and two when ignoring(response to the envelope modulation). For the group of musicians, we excluded one subject when attending and two when ignoring the speaker (response to the fundamental waveform). And lastly, one subject was excluded for both attending and ignoring the speaker, regarding the response to the envelope modulation. The subjects were excluded in all further analyses that only included the two musicality groups but were included in investigations on the whole subject cohort.

To assess which test we can use to compare the groups, for each combination of attention and frequency feature the distribution type of the musicians and non-musicians was computed. All distributions were normal except for the comparison between the two groups when attending to the speaker, regarding the response to the fundamental frequency. In this case, we used the Mann-Withney-U test whereas in the other cases, an unpaired Student’s t-test was applied. We found that there were no significant differences between musicians and non-musicians. We compared the two groups in all conditions for both mean and variance (Brown-Forsythe test).

**Figure 2:**
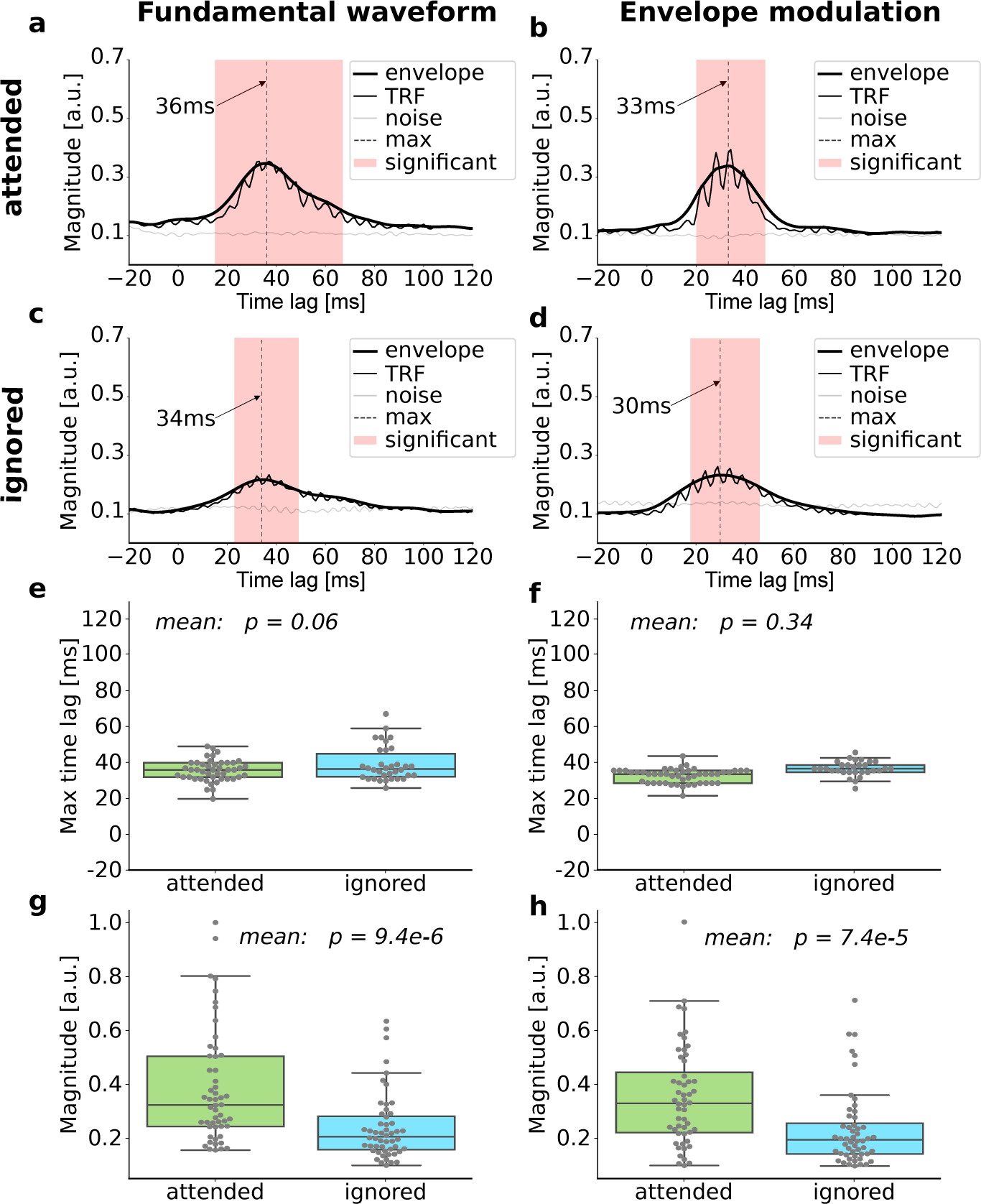
Attention modulation on the population level. (a)-(d) The amplitudes of the TRFs (thin black line) show significant activity at delays between about 20 ms and 50 ms (red shading) when compared to noise models (gray). They display some oscillatory activity at the fundamental frequency, which is no longer visible in the envelopes (thick black). The envelopes peak at latencies between 30 ms and 36 ms (vertical dashed line). (e,f) Subject-wise latencies of all significant envelope maxima (gray dots). They show no significant difference between attending and ignoring the speaker. The mean latencies of the distributions are used in the further analysis. (g,h) The magnitudes of the envelopes at the peak latencies are significantly higher in the attended than the ignored condition.

**Figure 3:**
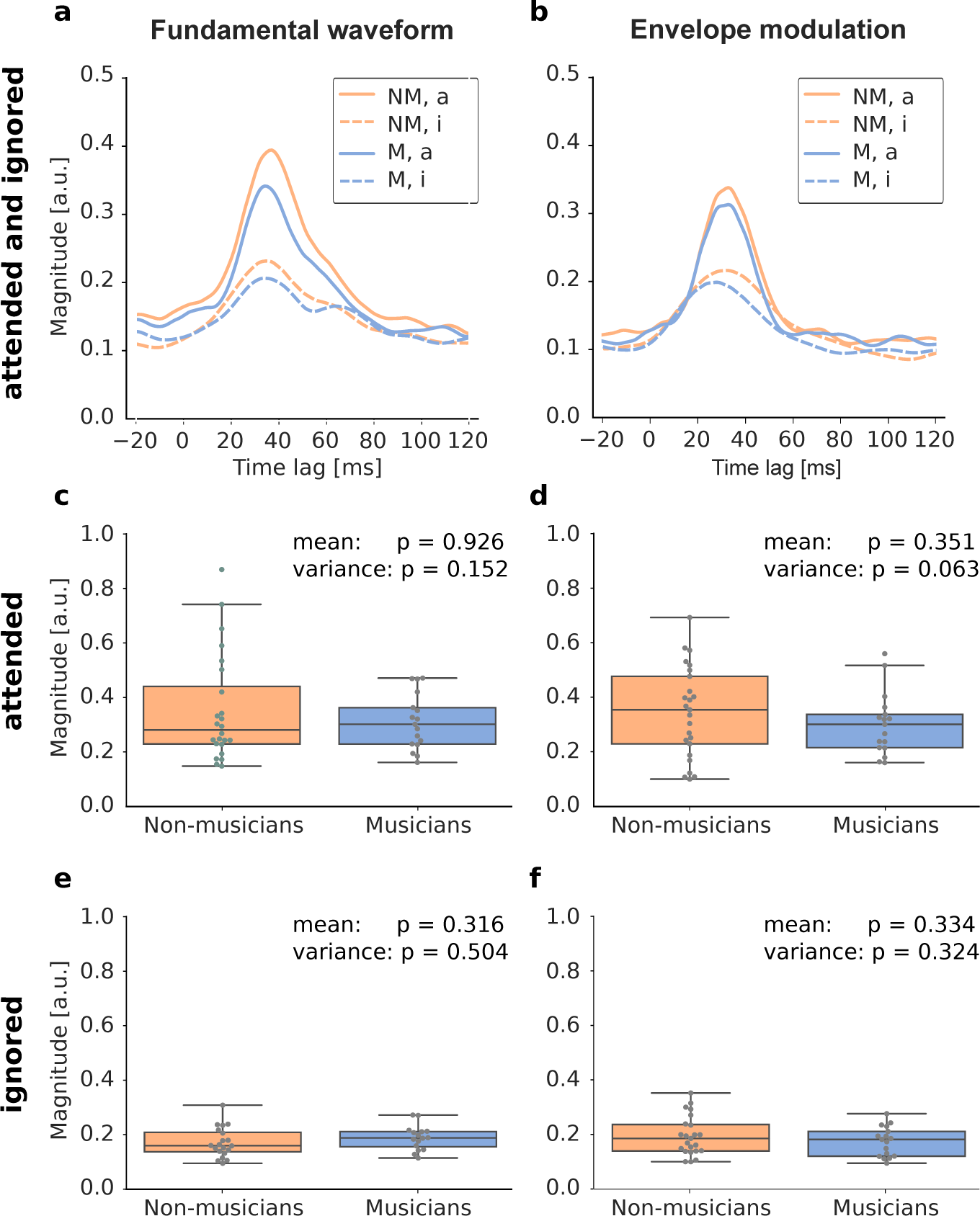
TRFs of musicians (M) versus non-musicians (NM). (a)-(b) Group-averaged normalized envelope TRFs for both non-musicians (blue) and musicians (orange) in the attended (a, solid) and ignored (i, dashed) condition for both acoustic features, respectively. (c)-(f) Comparison of the envelope values at the peak delays between the two groups, for both attention modes and for both acoustic features. The gray dots represent each subject’s magnitude. The *p*-values for the comparison of the means and of the variances between the two groups are included.

**Figure 4:**
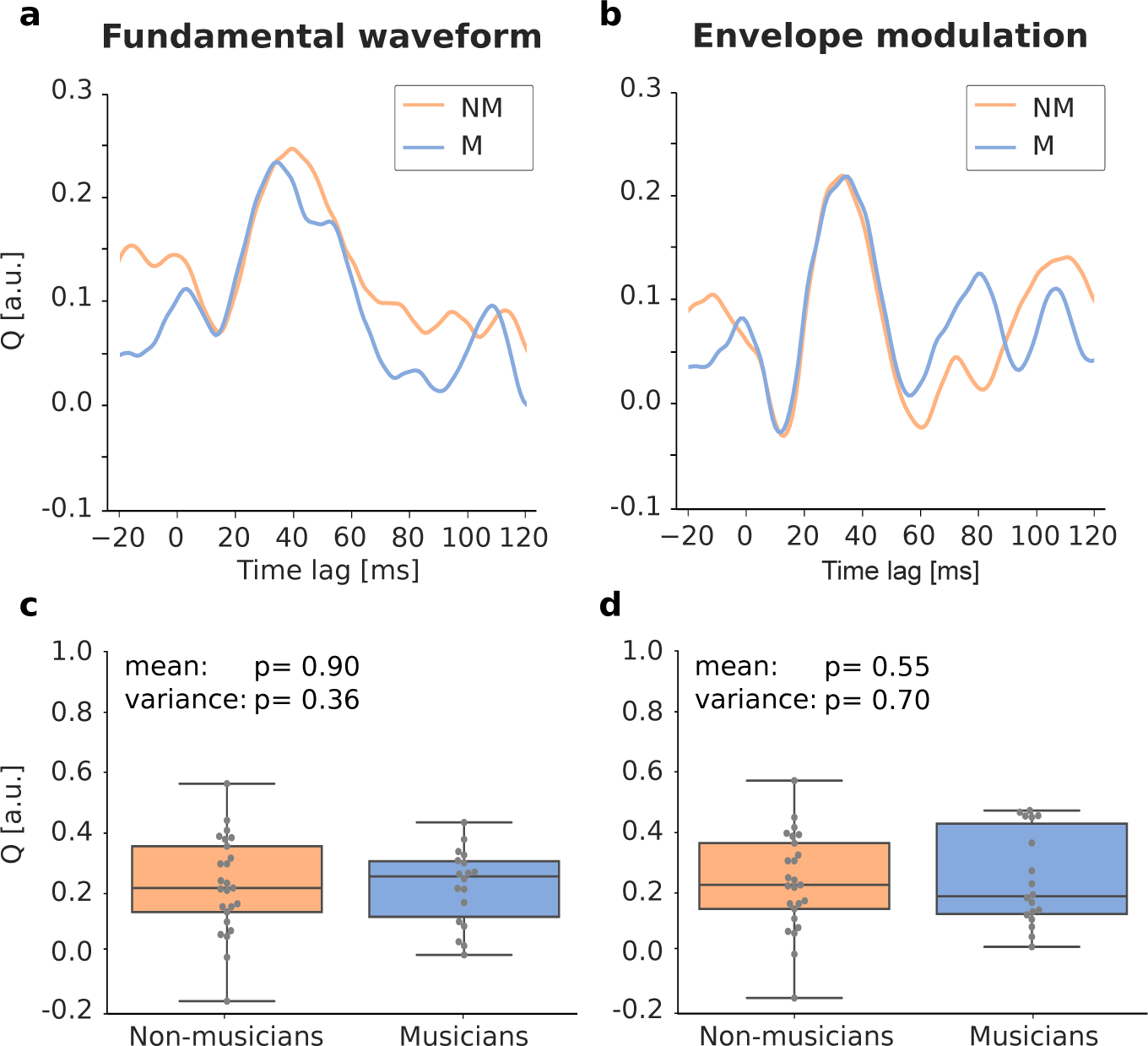
Attentional modulation in musicians (M) and non-musicians (NM). The effect of attention is quantified through the attentional modulation score *Q* computed from the envelopes of the TRF magnitudes. (a,b) The impact of attention on the brain signal is quantified by calculating the attentional modulation score *Q* for both musicians (M) and non-musicians (NM), at various latencies. (c,d) Investigating the subject-level values (gray dots) at the peak latencies did not reveal significant differences between musicians and nonmusicians.

### Comparison of the attentional modulation between musicians and non-musicians

To investigate the effect of musicality on the attentional modulation of the speech-FFR we calculated the difference quotient between attending and ignoring the speaker for both groups independently. This score *Q* is defined in Equation (2).

The attentional modulation score was computed separately for both musicians and non-musicians (Figure 4). To investigate systematic differences between the two groups, we focused on the values of the score *Q* per subject, at the peak latencies (Figure 4 (c,d)) and first computed their distribution type. We did, however, not find significant differences between musicians and non-musicians (fundamental waveform, mean: *p* = 0.90 unpaired t-test, variance: *p* = 0.36 Brown-Forsythe test; envelope modulation, mean: *p* = 0.55 Mann-Withney-U test, variance: *p* = 0.70 Brown-Forsythe test).

### Dependence of the cortical speech-FFR on the scores of musical training

In the previous analysis, we compared the group of musicians to that of non-musicians. This binary analysis ignored the variation and the different dimensions of musical training that the subjects received and also left some subjects out that fell in neither of the two groups. We therefore also investigated whether four different scores of musical training, as well as an aggregate score, could explain some of the subject-to-subject variation in the cortical speech-FFRs.

To this end, we once more considered the value of the envelope of the TRF magnitude at the peak latency, per subject. We then plotted these values against the five different scores and assessed whether a significant positive or negative correlation existed through the Spearman correlation coefficients (Figure 5). We considered all participants for this analysis, irrespective of whether they were classified as musicians, non-musicians, or neutral participants. The corresponding *p*-values after multiple comparison correction are also displayed in Figure 5. None of the correlations turned out to be statistically significant.

In Figure 5 (a) and (f), the neural responses are plotted against the age at which each subject first started training an instrument. For this analysis, the data from 15 non-musicians was not plotted as they never played an instrument and therefore did not have a starting age. In Figure 5 (b) and (g), the total number of years a participant had been practicing an instrument is displayed. Here, a very clear separation between the groups is visible (musicians: orange, non-musicians: blue, neither group: black).

In Figure 5 (c) and (h), the hours per week the participant presently trained an instrument are investigated. We note that one musician trained much more than the other participants, for more than 20 h/week.

The last score originating from the questionnaires is presented in Figure 5 (d) and (i) and displays the number of instruments the participant ever played in his or her life. As this criterion was not used for the grouping of the subjects, some mixture between the groups is visible. In Figure 5 (e) and (j), the neural response is plotted against the overall score.

### Dependence of the cortical speech-FFRs on the comprehension of the content questions

We wondered whether the performance of the participants in answering the comprehension question during the experiment was related to the neural responses. After each audiobook chapter, the subjects were asked to answer three multiple-choice questions with four possible answers, resulting in a chance level of 25%. The percentage of questions answered correctly and the corresponding value of the cortical speech-FFR for each participant, obtained from the envelope of the TRF magnitude at the peak latency, is presented in Figure 6 for both acoustic features.

Computation of Spearman’s correlation coefficients and testing of their statistical significance shows the absence of any significant correlation (fundamental waveform, *r* = *−*0.02, *p* = 0.87; envelope modulation, *r* = *−*0.14, *p* = 0.33).

Furthermore, the behavioral results were compared between the three groups of participants, independent of their neural responses. Therefore, the number of correctly answered questions of each participant was used and the Kruskal test was applied to compare the three groups. The test provided a *p*-value of 0.64, meaning there was no significant difference between the groups.

**Figure 5:**
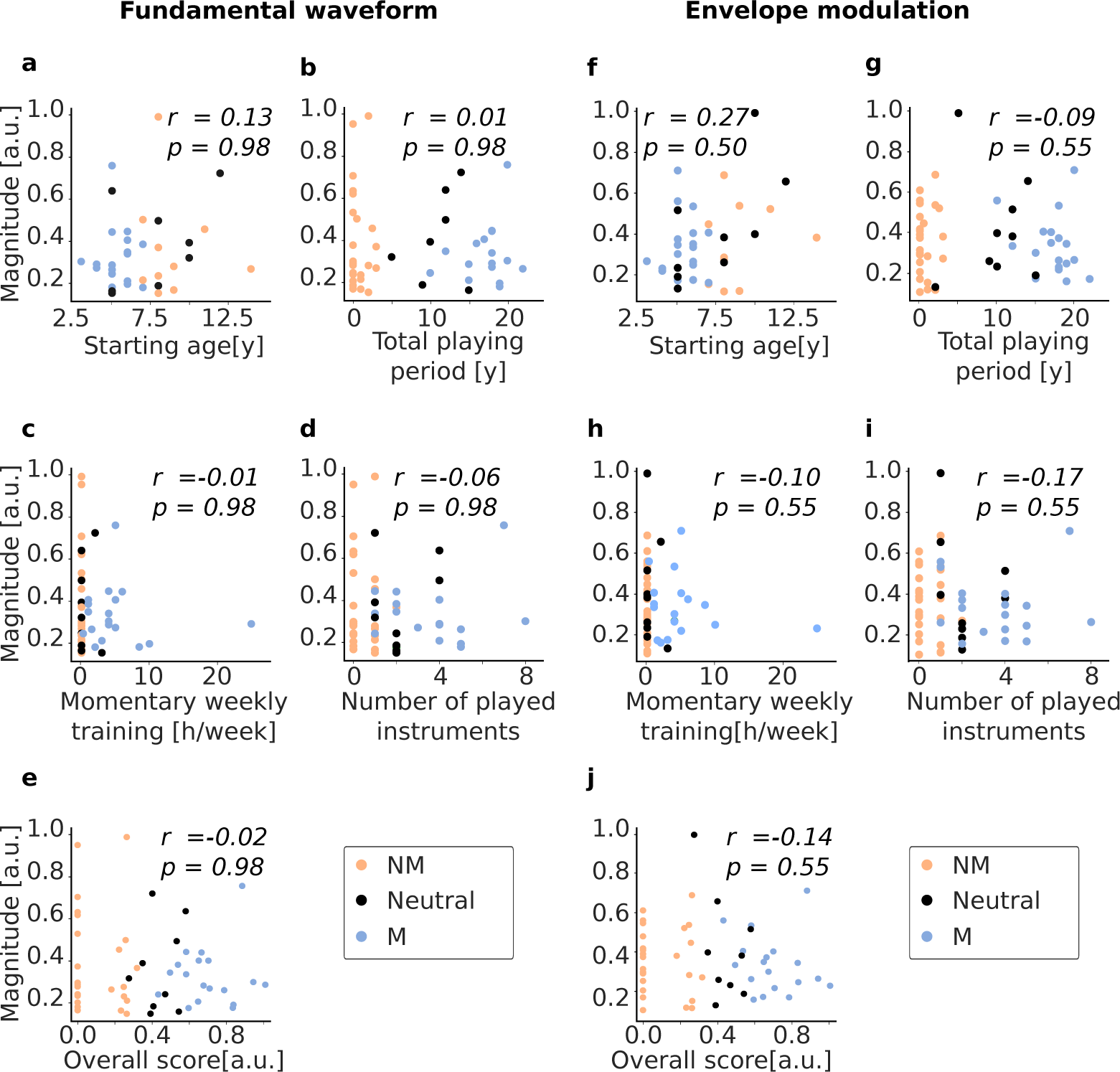
Dependence of the magnitude of the cortical speech-FFRs on four scores of musical training as well as on an aggregate score. The results for the fundamental waveform are plotted in (a)-(e) and for the envelope modulation in (f)-(j). Non-musicians (NM) are displayed in orange, musicians (M) in blue, and participants belonging to neither group (neutral) in black. None of the dependencies is statistically significantly correlated.

**Figure 6:**
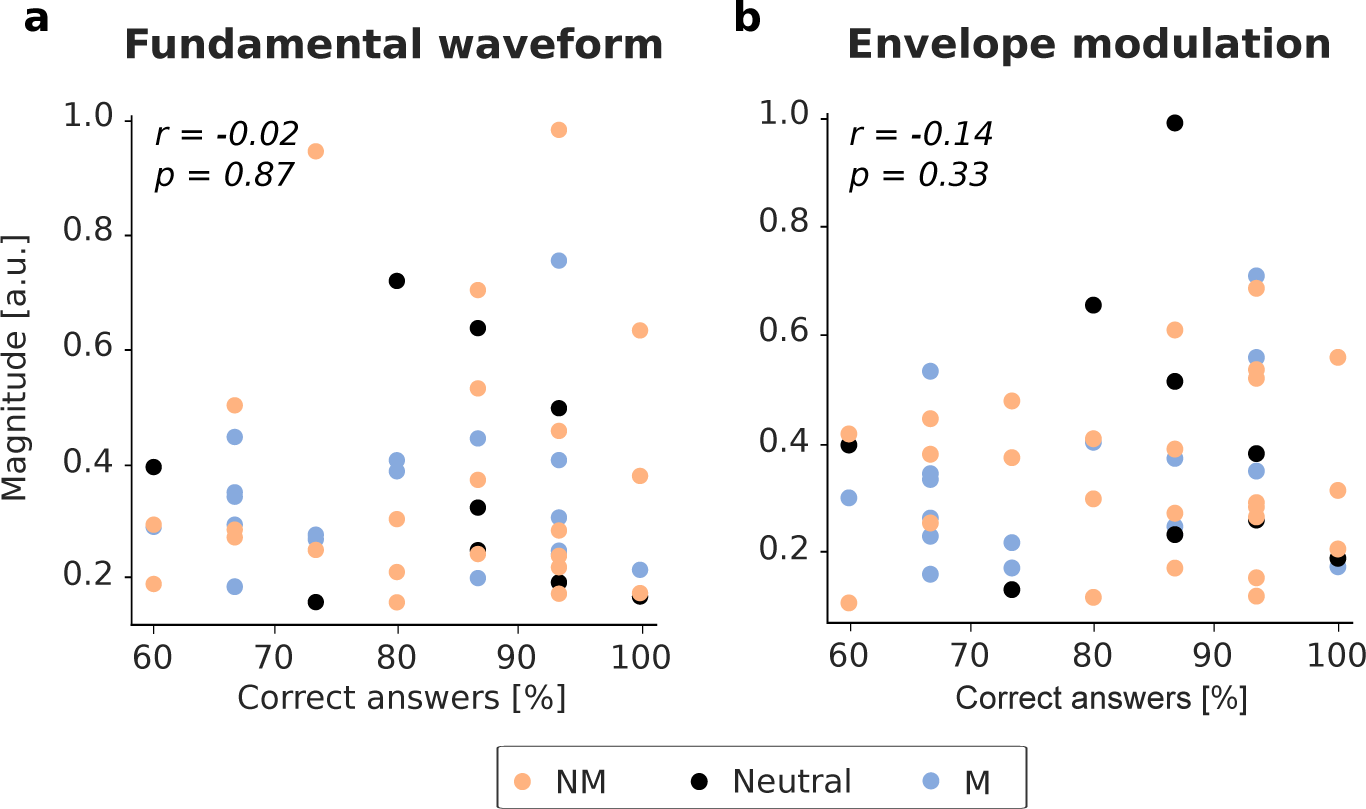
Relation between the cortical speech-FFR and the comprehension of the content questions for individual subjects. Neither the neural responses to the fundamental waveform (a) nor those to the envelope modulation (b) showed a statistically significant dependence.

## Discussion

This study is the first to examine the influence of musical training on the cortical contribution to the speech-FFR, as well as on its attentional modulation. To this end, we employed MEG recordings of participants who were asked to attend one of two continuous competing speech streams. The cortical speech-FFRs were analyzed through source reconstruction followed by the computation of temporal response functions (TRFs). The TRFs related the neural data in cortical regions of interest to two features of the speech signal that were related to the fundamental frequency: the fundamental waveform and the envelope modulation.

### Measurement of cortical speech-FFRs and their attentional modulation

The cortical contribution to the speech-FFR was only discovered a few years ago, in addition to the well-known subcortical contribution that occurs at a delay of around 10 ms (Coffey et al., 2016, 2019; Kulasingham et al., 2020). The observation of its attentional modulation is even more recent (Schüller et al., 2023a; Commuri et al., 2023). Our measurements in a large group of participants reported here confirm that these cortical contributions can be measured reliably. We observed maximal responses at time lags between 30 ms and 36 ms, in agreement with the previous measurements of the cortical contribution to the speech-FFR (Kulasingham et al., 2020; Schüller et al., 2023b). This supports the cortical origin of the signals that we consider here.

In addition, our study confirmed a sizable effect of attention on the cortical speech-FFRs, with the neural responses being significantly larger when the speaker was attended than when he was ignored. The origin of these attention effects is still unclear: they might merely reflect attentional effects on the subcortical contribution to the speech-FFR, or attentional modulation due to top-down feedback from higher areas of the auditory cortex (Forte et al., 2017; Etard et al., 2019; Saiz-Aĺıa et al., 2019).

Regarding the two speech features that we employed, our study found that the signals were of similar strength for the envelope modulation compared to the fundamental waveform, regardless of whether they were attended or ignored. Previous research conducted by Kulasingham et al. (2020) and Schüller et al. (2023a) using a similar experimental setup showed enhanced TRF strength for envelope modulation. Additionally, Kegler et al. (2022a) reported similar results in the subcortical region when examining word-level acoustic and linguistic information as input signals using EEG.

### Differences in the cortical speech-FFRs between musicians and non-musicians

To investigate the influence of musical training, we first divided our participants into three groups: musicians, non-musicians, and those who fell into neither of the two former groups. Surprisingly, musicians and non-musicians exhibited comparable cortical speech-FFRs. In particular, both mean and variance for both attending and ignoring the speaker, evaluated for fundamental waveform and envelope modulation were not significant. This suggests that musical training does not affect the cortical contribution to the speech-FFR.

This result is surprising as the subcortical contribution to the speech-FFR, as well as other subcortical FFRs, are well-known to be larger in musicians than in non-musicians (Musacchia et al., 2008a; Weiss and Bidelman, 2015; Bidelman et al., 2014; Musacchia et al., 2008b; Parbery-Clark et al., 2012a; Rodrigues et al., 2019). The enhanced subcortical speech-FFR in people with musical training is assumed to explain enhanced behavioral performance in musicians, such as better speech-in-noise comprehension (Parbery-Clark et al., 2009, 2012a) and better pitch discrimination Magne et al. (2006).

Our finding that musicians exhibit cortical speech-FFRs with similar strength as non-musicians suggests that the cortical contribution to the speech-FFR does not merely reflect the corresponding subcortical activity, but is influenced by additional processing. This viewpoint is corroborated by the observation that the cortical contribution to the speech-FFR is right-lateralized, which is not the case for the subcortical sources (Coffey et al., 2016; Schüller et al., 2023b). Effects on the cortical signal strength in musicians have so far only been found for input signals like complex tones (Bianchi et al., 2016). These findings might relate more to musical experiences and less to how speech is processed.

As we did not find any significant differences between both groups we wondered if our criteria for defining which subject is a musician might not be strict enough for them to show differences. We did indeed not observe behavioral differences between the different groups; all answered the comprehension questions that we posed equally well. We note, however, that we did not design the comprehension questions for a fine-grained measure of speech-in-noise ability, but merely to verify attention to the target speaker. Moreover, we employed criteria to categorize the participants as musicians/non-musicians that are the same or very similar to those of studies that did find differences. Moreover, several studies show effects of musical training on a continuous scale (Strait et al., 2009; Pantev et al., 1998; Musacchia et al., 2007). This would not be the case if a specific threshold in musical training had to be reached.

### Influence of musical training on the attentional modulation

The modulation of the cortical speech-FFRs by selective attention to one of the two competing speakers may aid speech-in-noise comprehension, by aiding an enhanced neural representation of the target speech. We, therefore, hypothesized that the better speech-in-noise comprehension abilities of musicians may originate in a larger attentional modulation of the cortical speech-FFRs. However, when comparing the latter between musicians and non-musicians, we did not find a significant difference.

This result implies that the larger cortical speech-FFRs and their larger variation observed in non-musicians are not attributable to the influence of musical training on the attentional effect.

There have only been a few prior studies on musical training and attentional modulation of neural responses. In particular, it has not yet been investigated whether the attentional modulation of the subcortical speech-FFR is influenced by musical training. The only studies on this issue investigated event-related potentials (ERPs) and found either no significant differences Tervaniemi et al. (2004) or enhanced attentional effects Tervaniemi et al. (2009). Significant additional research is required to more fully investigate how musical training shapes the responses along the auditory pathway, from the brainstem through the midbrain and the various processing centers of the auditory cortex.

### Relationships between different aspects of musical training on the cortical speech-FFR

Besides comparing musicians to non-musicians, we also investigated whether the cortical speech-FFRs depended on specific aspects of musical training. This analysis included all subjects, including those who did not qualify as either musicians or non-musicians.

The scores of musical training included the age at which participants started playing their first instrument (if applicable), the total number of years participants had been playing any type of instrument, the amount of training the participant was currently undertaking, and the number of instruments the participant had experience playing. Additionally, a fifth score was calculated as an aggregate of the four scores.

When computing Spearman’s correlation coefficients between the cortical speech-FFRs and the different scores of musical training, we found none that were statistically significant. As presented above, this is in contrast to the subcortical contribution to the speech-FFR. For instance, in a study by Strait et al. (2009), it was demonstrated that the amplitude of the subcortical response was positively correlated with the number of years of consistent practice. Furthermore, the subcortical contribution was larger for individuals who began their musical training at a younger age (Strait et al., 2009).

### Relation of the subcortical to the cortical contributions to the speech-FFR

There has not yet been a combined EEG and MEG study measuring the cortical and subcortical contribution to the speech-FFR in parallel. However, a recent publication showed that subcortical and cortical contributions to the speech-FFR can be measured simultaneously in the same MEG dataset (Schüller et al., 2023b). Both could well be separated based on their delays: the subcortical response occurred at a delay of around 10 ms, whereas the cortical response, similar to the ones reported here, exhibited latencies of around 35 ms. However, due to the poor sensitivity of MEG to subcortical sources, the subcortical responses were weak and could only be measured due to the long recording times and the noise-free audio signal with a single speaker.

In the dataset that we analyzed here, with two competing speakers and alternating attention, we were unfortunately not able to identify the subcortical contribution. However, since prior EEG studies showed that the subcortical responses are modulated by attention as well (Forte et al., 2017), attention may likely impact the cortical and subcortical responses through partly shared efferent pathways. How musical training affects these top-down effects may help to further elucidate these neural feedback loops.

## Conclusion

In summary, we have conducted the first study that investigated the impact of musical training on the cortical contribution to the speech-FFR. Contrary to the subcortical contribution, where musical training leads to enhanced neural responses, the cortical contribution was similar for musicians and non-musicians. The attentional modulation of the cortical speech-FFR was not influenced by musical training either. Moreover, we did not find that specific scores of musical training were correlated to the strength of the cortical speech-FFR. Taken together, our results show that musical training has a weakly detrimental effect on the cortical response. This evidences that the subcortical and the cortical contribution to the speech-FFR play at least partly different roles in the processing of complex acoustic stimuli such as speech and music.

## References

Benjamini Y, Hochberg Y (1995) Controlling the false discovery rate: A practical and powerful approach to multiple testing. Journal of the Royal Statistical Society: Series B (Methodological) 57:289–300.

Besson M, Schön D, Moreno S, Santos A, Magne C (2007) Influence of musical expertise and musical training on pitch processing in music and language. Restor. Neurol. Neurosci. 25:399–410.

Bianchi F, Santurette S, Wendt D, Dau T (2016) Pitch discrimination in musicians and non-musicians: Effects of harmonic resolvability and processing effort. Journal of the Association for Research in Otolaryngology 17:69–79.

Bidelman GM, Gandour JT, Krishnan A (2011) Musicians and tone-language speakers share enhanced brainstem encoding but not perceptual benefits for musical pitch. Brain and Cognition 77:1–10.

Bidelman GM, Weiss MW, Moreno S, Alain C (2014) Coordinated plasticity in brainstem and auditory cortex contributes to enhanced categorical speech perception in musicians. European Journal of Neuroscience 40:2662–2673.

Bourgeois J, Minker W (2009) Linearly constrained minimum variance beamforming. Time-Domain Beam-forming and Blind Source Separation: Speech Input in the Car Environment pp. 27–38.

Bronkhorst AW (2000) The cocktail party phenomenon: A review of research on speech intelligibility in multiple-talker conditions. Acta Acustica united with Acustica 86:117–128.

Chi T, Ru P, Shamma SA (2005) Multiresolution spectrotemporal analysis of complex sounds. The Journal of the Acoustical Society of America 118:887–906.

Coffey EBJ, Herholz SC, Chepesiuk AMP, Baillet S, Zatorre RJ (2016) Cortical contributions to the auditory frequency-following response revealed by MEG. Nature Communications 7.

Coffey EB, Nicol T, White-Schwoch T, Chandrasekaran B, Krizman J, Skoe E, Zatorre RJ, Kraus N (2019) Evolving perspectives on the sources of the frequency-following response. Nature communications 10:5036.

Commuri V, Kulasingham JP, Simon JZ (2023) Cortical responses time-locked to continuous speech in the high-gamma band depend on selective attention. Frontiers in Neuroscience 17.

Deroche MLD, Limb CJ, Chatterjee M, Gracco VL (2017) Similar abilities of musicians and non-musicians to segregate voices by fundamental frequency. The Journal of the Acoustical Society of America 142:1739–1755.

Douw L, Nieboer D, Stam CJ, Tewarie P, Hillebrand A (2018) Consistency of magnetoencephalographic functional connectivity and network reconstruction using a template versus native mri for co-registration. Human Brain Mapping 39:104–119.

Eaves JM, Quentin Summerfield A, Kitterick PT (2011) Benefit of temporal fine structure to speech perception in noise measured with controlled temporal envelopes. The Journal of the Acoustical Society of America 130:501–507.

Etard O, Kegler M, Braiman C, Forte AE, Reichenbach T (2019) Decoding of selective attention to continuous speech from the human auditory brainstem response. NeuroImage 200:1–11.

Fischl B (2012) Freesurfer. NeuroImage 62:774–781.

Forte AE, Etard O, Reichenbach T (2017) The human auditory brainstem response to running speech reveals a subcortical mechanism for selective attention. eLife 6.

Gramfort A, Luessi M, Larson E, Engemann DA, Strohmeier D, Brodbeck C, Parkkonen L, Hämäläinen MS (2014) MNE software for processing MEG and EEG data. NeuroImage 86:446–460.

Holliday IE, Barnes GR, Hillebrand A, Singh KD (2003) Accuracy and applications of group meg studies using cortical source locations estimated from participants’ scalp surfaces. Human Brain Mapping 20:142–147.

Hopkins K, Moore BC (2009) The contribution of temporal fine structure to the intelligibility of speech in steady and modulated noise. The Journal of the Acoustical Society of America 125:442–446.

Kaplan EC, Wagner AE, Toffanin P, Başkent D (2021) Do musicians and non-musicians differ in speech-on-speech processing? Frontiers in Psychology 12.

Kaptein M, van den Heuvel E (2022) Statistics for Data Scientists Springer International Publishing.

Kegler M, Weissbart H, Reichenbach T (2022a) The neural response at the fundamental frequency of speech is modulated by word-level acoustic and linguistic information. Frontiers in Neuroscience 16.

Kegler M, Weissbart H, Reichenbach T (2022b) The neural response at the fundamental frequency of speech is modulated by word-level acoustic and linguistic information. Frontiers in Neuroscience 16.

Kraus N, Chandrasekaran B (2010) Music training for the development of auditory skills. Nature Reviews Neuroscience 11:599–605.

Kulasingham JP, Brodbeck C, Presacco A, Kuchinsky SE, Anderson S, Simon JZ (2020) High gamma cortical processing of continuous speech in younger and older listeners. NeuroImage 222:117291.

Madsen SMK, Marschall M, Dau T, Oxenham AJ (2019) Speech perception is similar for musicians and non-musicians across a wide range of conditions. Scientific Reports 9.

Magne C, Schön D, Besson M (2006) Musician children detect pitch violations in both music and language better than nonmusician children: Behavioral and electrophysiological approaches. Journal of Cognitive Neuroscience 18:199–211.

Maillard E, Joyal M, Murray MM, Tremblay P (2023) Are musical activities associated with enhanced speech perception in noise in adults? a systematic review and meta-analysis. Current Research in Neurobiology 4:100083.

Mauch M, Dixon S (2014) pYIN: A fundamental frequency estimator using probabilistic threshold distributions In 2014 IEEE International Conference on Acoustics, Speech and Signal Processing (ICASSP), pp. 659–663. IEEE.

McDermott JH (2009) The cocktail party problem. Current Biology 19:R1024–R1027.

Meha-Bettison K, Sharma M, Ibrahim RK, Vasuki PRM (2017) Enhanced speech perception in noise and cortical auditory evoked potentials in professional musicians. International Journal of Audiology 57:40–52.

Morse-Fortier C, Parrish MM, Baran JA, Freyman RL (2017) The effects of musical training on speech detection in the presence of informational and energetic masking. Trends in Hearing 21:233121651773942.

Musacchia G, Sams M, Skoe E, Kraus N (2007) Musicians have enhanced subcortical auditory and audiovisual processing of speech and music. Proceedings of the National Academy of Sciences 104:15894–15898.

Musacchia G, Strait D, Kraus N (2008a) Relationships between behavior, brainstem and cortical encoding of seen and heard speech in musicians and non-musicians. Hearing research 241:34–42.

Musacchia G, Strait D, Kraus N (2008b) Relationships between behavior, brainstem and cortical encoding of seen and heard speech in musicians and non-musicians. Hearing Research 241:34–42.

Pantev C, Oostenveld R, Engelien A, Ross B, Roberts LE, Hoke M (1998) Increased auditory cortical representation in musicians. Nature 392:811–814.

Pantev C, Roberts LE, Schulz M, Engelien A, Ross B (2001) Timbre-specific enhancement of auditory cortical representations in musicians. Neuroreport 12:169–174.

Parbery-Clark A, Tierney A, Strait D, Kraus N (2012a) Musicians have fine-tuned neural distinction of speech syllables. Neuroscience 219:111–119.

Parbery-Clark A, Anderson S, Hittner E, Kraus N (2012b) Musical experience strengthens the neural representation of sounds important for communication in middle-aged adults. Frontiers in Aging Neuro-science 4.

Parbery-Clark A, Skoe E, Kraus N (2009) Musical experience limits the degradative effects of background noise on the neural processing of sound. The Journal of Neuroscience 29:14100–14107.

Parbery-Clark A, Skoe E, Lam C, Kraus N (2009) Musician enhancement for speech-in-noise. Ear &amp Hearing 30:653–661.

Perrot X, Micheyl C, Khalfa S, Collet L (1999) Stronger bilateral efferent influences on cochlear biome-chanical activity in musicians than in non-musicians. Neuroscience Letters 262:167–170.

Rodrigues M, Donadon C, Guedes-Weber M, Sant’anna S, Skarzynski P, Hatzopoulos S, Colella-Santos M, Sanfins M (2019) Frequency following resonse and musical experience: a review. Journal of Hearing Science 9:9–16.

Russo N, Nicol T, Musacchia G, Kraus N (2004) Brainstem responses to speech syllables. Clinical Neuro-physiology 115:2021–2030.

Saiz-Alía M, Forte AE, Reichenbach T (2019) Individual differences in the attentional modulation of the human auditory brainstem response to speech inform on speech-in-noise deficits. Scientific Reports 9.

Saiz-Alía M, Reichenbach T (2020) Computational modeling of the auditory brainstem response to continuous speech. Journal of Neural Engineering 17:036035.

Santos A, Oivo M, Juristo N (2018) Moving beyond the mean: Analyzing variance in software engineering experiments In Product-Focused Software Process Improvement, pp. 167–181. Springer International Publishing.

Schilling A, Tomasello R, Henningsen-Schomers MR, Zankl A, Surendra K, Haller M, Karl V, Uhrig P, Maier A, Krauss P (2021) Analysis of continuous neuronal activity evoked by natural speech with computational corpus linguistics methods. *Language*, Cognition and Neuroscience 36:167–186.

Schüller A, Schilling A, Krauss P, Rampp S, Reichenbach T (2023a) Attentional modulation of the cortical contribution to the frequency-following response evoked by continuous speech. Journal of Neuroscience 43:7429–7440.

Schüller A, Schilling A, Krauss P, Reichenbach T (2023b) The early subcortical response at the fundamental frequency of speech is temporally separated from later cortical contributions. Journal of Cognitive Neuroscience pp. 1–17.

Skoe E, Kraus N (2010) Auditory brainstem response to complex sounds: a tutorial. Ear and hearing 31:302.

Strait DL, Kraus N, Skoe E, Ashley R (2009) Musical experience and neural efficiency-effects of training on subcortical processing of vocal expressions of emotion. European Journal of Neuroscience 29:661–668.

Tervaniemi M, Kruck S, Baene WD, Schröger E, Alter K, Friederici AD (2009) Top-down modulation of auditory processing: effects of sound context, musical expertise and attentional focus. European Journal of Neuroscience 30:1636–1642.

Tervaniemi M, Just V, Koelsch S, Widmann A, Schrger E (2004) Pitch discrimination accuracy in musicians vs nonmusicians: an event-related potential and behavioral study. Experimental Brain Research 161:1–10.

Toh XR, Tan SH, Wong G, Lau F, Wong FC (2023) Enduring musician advantage among former musicians in prosodic pitch perception. Scientific Reports 13:2657.

Van Canneyt J, Wouters J, Francart T (2021) Neural tracking of the fundamental frequency of the voice: the effect of voice characteristics. European Journal of Neuroscience 53:3640–3653.

van Zuijen TL, Sussman E, Winkler I, Näätänen R, Tervaniemi M (2005) Auditory organization of sound sequences by a temporal or numerical regularity—a mismatch negativity study comparing musicians and non-musicians. Cognitive Brain Research 23:270–276.

Watanabe D, Savion-Lemieux T, Penhune VB (2006) The effect of early musical training on adult motor performance: evidence for a sensitive period in motor learning. Experimental Brain Research 176:332–340.

Weiss MW, Bidelman GM (2015) Listening to the brainstem: musicianship enhances intelligibility of subcortical representations for speech. Journal of Neuroscience 35:1687–1691.

Wong DD, Fuglsang SA, Hjortkjær J, Ceolini E, Slaney M, de Cheveigné A (2018) A comparison of temporal response function estimation methods for auditory attention decoding. *bioRxiv*.

Wong PCM, Skoe E, Russo NM, Dees T, Kraus N (2007) Musical experience shapes human brainstem encoding of linguistic pitch patterns. Nature Neuroscience 10:420–422.

Zhang L, Fu X, Luo D, Xing L, Du Y (2020) Musical experience offsets age-related decline in understanding speech-in-noise: Type of training does not matter, working memory is the key. Ear amp; Hearing 42:258–270.

